# Alpha-2-Macroglobulin mitigates mitochondrial dysfunction and neuronal apoptotic responses in pesticide-induced neurotoxicity

**DOI:** 10.1101/2025.06.27.661925

**Authors:** Swati Dixit

## Abstract

Persistent and repetitive application of pesticides has been linked to adverse effects on human metabolism and the onset of various disorders. Commonly used pesticides, such as carbamates (e.g., Aldicarb, ALD) and organophosphates (e.g., Chlorpyrifos, CPF), are widely applied in potato cultivation and household pest control. Chronic exposure to these substances has been implicated in the early onset of neurodegenerative disorders. Alpha-2-macroglobulin (A2M), a protein known for its regulatory role in oxidative stress, participates in multiple biological processes. Despite its significance, the role of A2M in mitigating mitochondrial-induced neuronal apoptosis triggered by pesticide interference remains poorly understood. This study explores the involvement of A2M in SH-SY5Y neuroblastoma cells exposed to pesticides, focusing on its impact on mitochondrial enzyme expression, inflammatory cytokines, neuronal apoptotic and anti-apoptotic proteins, and the activation of the Nrf2 signaling pathway. Comparative analyses of control and pesticide-exposed SH-SY5Y cells revealed that A2M positively modulates neuronal stress responses. Western blot profiling demonstrated that A2M upregulates anti-apoptotic proteins such as Bcl-2 and Nrf2, while downregulating pro-apoptotic markers, including Bax, Caspase-3, and Caspase-9. Biochemical assays showed that A2M enhances mitochondrial enzyme activity, particularly complexes I and III, while mitigating the reactive oxygen species (ROS) generated by ALD and CPF exposure. Furthermore, A2M was found to reduce DNA damage caused by pro-inflammatory cytokines, which are exacerbated by mitochondrial oxidative stress. These findings highlight the pivotal role of A2M in attenuating pesticide-induced neuronal toxicity through the regulation of mitochondrial function and inhibition of neuronal apoptosis.

**Research Highlights:**

- This study is the first to report the interaction between human alpha-2-macroglobulin and neuronal apoptotic markers in pesticides mediated neurotoxicity.
- The study found that treatment of alpha-2-macroglobulin promotes the proliferation of Nrf2 pathway during oxidative stress in neuronal cells.
- This research reveals that alpha-2-macroglobulin modulates oxidative stress in mitochondrial enzyme complexes and during mitochondrial dysfunction.
- The study confirms that treatment of alpha-2-macroglobulin significantly alleviates pro inflammatory cytokines during pesticides mediated neurotoxicity.
- This research provides new theoretical basis and molecular targets for the treatment of neurodegenerative diseases during early preclinical stages, offering broad application prospects.

**GRAPHICAL ABSTRACT:** 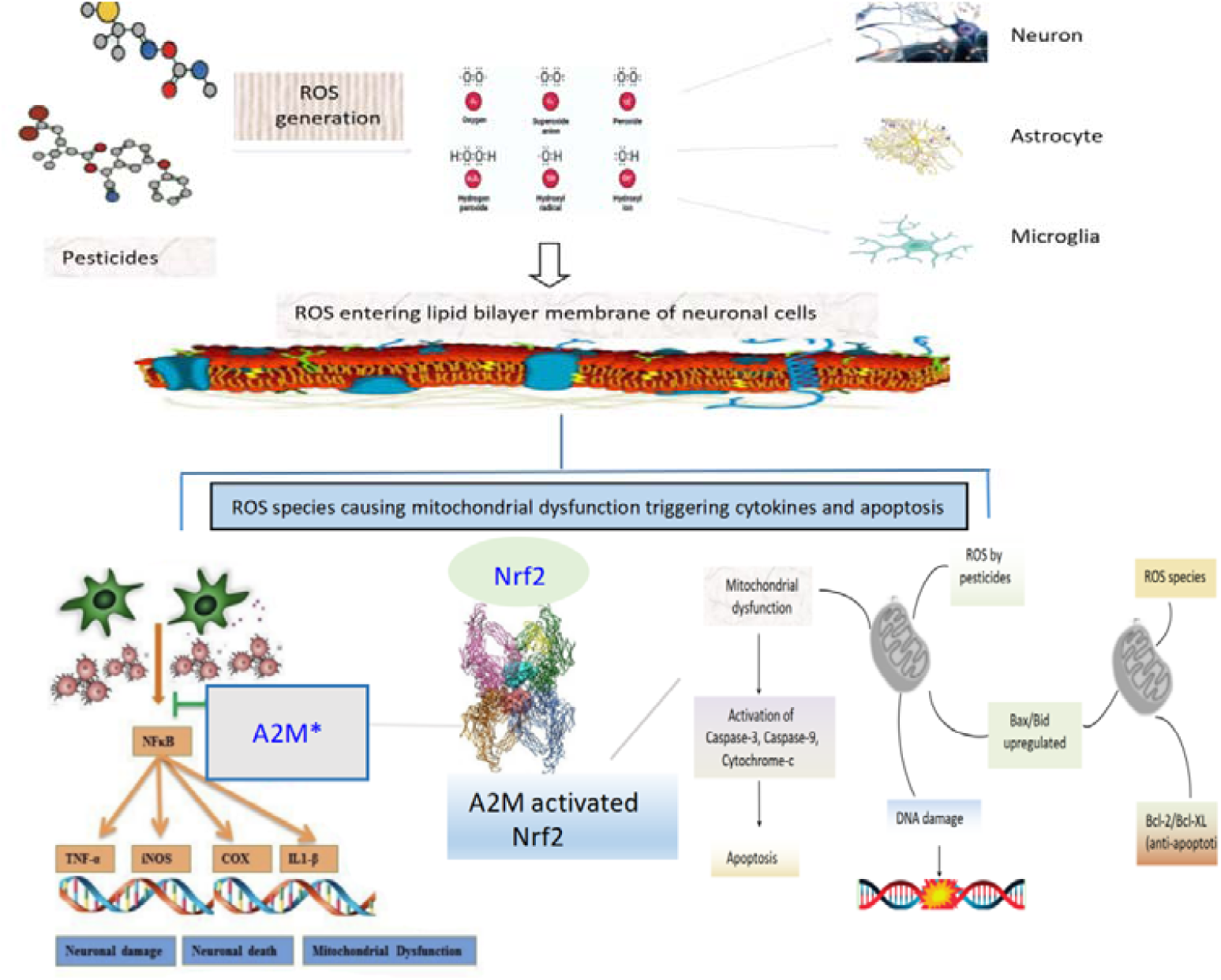

## INTRODUCTION

Neurodegenerative diseases such as Alzheimer’s disease (AD), Parkinson’s disease (PD), Multiple sclerosis (MS) are major clinical concern worldwide. Epidemiological evidences reports dietary, genetic, molecular and environmental factors are leading factors of neurodegeneration (**Chin-Chan et al., 2015**). The contaminants accumulate in human body via food chain and environmental factors, thus altering gene expression and modify protein synthesis and significantly affect other biological process (**Yu et al., 2021a**). Pesticides such as carbamates, pyrethroids, organophosphates are recognized as one of the risk factors involved in epigenetic mechanisms leading to progression of neurodegenerative diseases (**Yu et al., 2021b**). Low levels of pesticides accumulate in tissues, impact human brain causing impairment in cognitive functions, memory and some motor functions (**Paul et al., 2018**). These neurobehavioral changes are associated with the development and pathogenesis of various neurological disorders (**Dixit et al., 2023**).

One of the prominent neurodegenerative disease worldwide is Alzheimer’s disease, which is characterised by loss of neurons and synapses in the cerebral cortex and certain subcortical regions (**Varma et al., 2017**). Preclinical stages of AD can be determined by examining the Cererbrospinal fluid (CSF), neuroimaging of biomarkers to recognise the deposition of β-amyloid plaques, synaptic dysfunction and inflammatory and apoptotic markers of neuronal injury (**Dubois et al., 2014**). Some inflammatory responses and immune regulatory factors are intrinsic to AD pathogenesis and act as trigger of AD onset and its clinical symptoms (**Heneka et al., 2015**). These factors include pro inflammatory cytokines, apoptotic protein markers, transcription factors etc. (**Chen et al., 2024**). Research suggests that exposure to certain pesticides interferes with normal immune system functioning, leading to increased inflammation (**Li et al., 2022**). These chemicals may interfere with the functioning of immune cells, such as macrophages, by altering their cytokines production and can cause mitochondrial dysfunction in neurons which further lead to alterations in DNA expression and triggers apoptosis (**Alimova et al., 2022**).

Alpha-2 macroglobulin (A2M) is an acute phase protein and major component of the innate immune system, functioning as a pan-protease inhibitor and a chaperone protein (**French et al., 2008, Varma et al., 2017, Dixit et al., 2021a**). A2M is a tetramer, consisting of bait region located among the each identical subunits which functionally inhibits any proteinases present in plasma regardless of their specificity (**fig.1**)(**Dixit et al., 2021a**). It transports and inhibits pro inflammatory cytokines such as IL-1β, IL-6, interferons, insulin, growth hormone, and transforming growth factor-β and tumor necrosis factor (TNF-α), platelet derived growth factor etc. (**Dixit et al., 2021b**). Affinity of A2M to cytokines has led to efforts targeting inflammation as a potential strategy for AD pathogenesis. Although, the strategy of targeting anti-inflammatory agents have failed in the patients diagnosed with established AD have failed, which alternatively suggests targeting inflammation and anti-inflammatory substances during early stage before the onset of clinical symptoms of AD (**McCaulley and Grush, 2015**).

**Figure 1:**
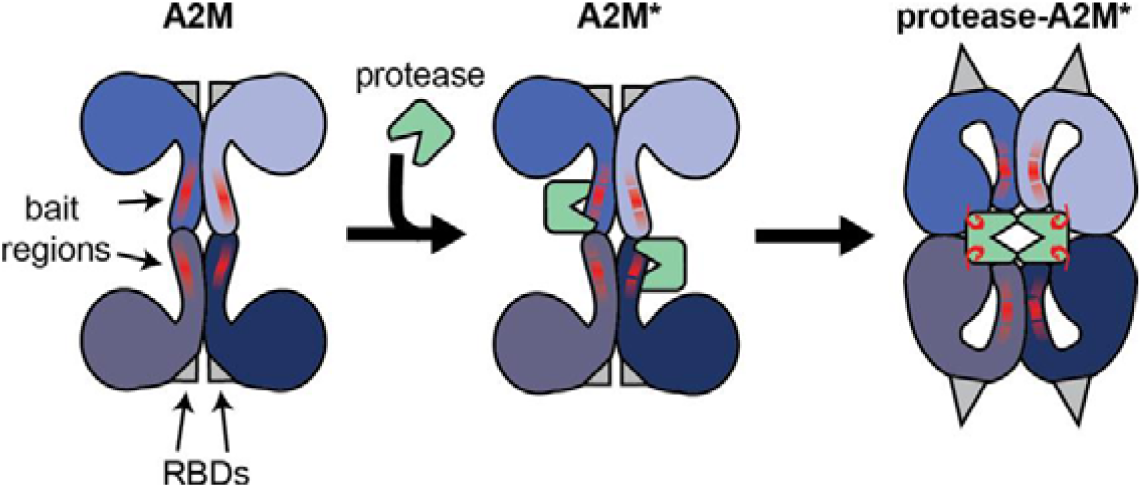
Representation of the interaction between the A2M and proteases. Each subunit contains a protease bait region and a buried receptor binding domain. Proteases recognise the bait region and interacts with it, which changes the protein’s conformation resulting in trapping the proteases.

In our previous study, we have investigated the cytoprotective role of A2M against pesticides (Chlorpyrifos, Aldicarb and Deltamethrin) induced neurotoxicity targeting its role in modulating antioxidant enzymes and ROS species markers such as MDA, TBARS in neuronal SH-SY5Y cell lines. We found that A2M shielded the antioxidant system of cells against ROS (**Dixit et al., 2023**), however some other issues need to be addressed to reach some conclusion about A2M’s protective mechanism (**Dixit et al., 2022b**). This present study evaluates the neuroprotective effects of A2M on pesticides induced mitochondrial dysfunction, inflammatory aberrations, and neuronal apoptosis to identify A2M’s ameliorative effective as a strategic approach to early inflammatory and immune-regulatory processes which are intrinsic to AD pathogenesis. Our study focuses whether A2M renders neuroprotection by inhibiting mitochondrial mediated apoptotic pathway and underlying events like apoptosis in neuronal SH-SY5Y cells during pesticides induced neurotoxicity; whether A2M reduces neuronal inflammation by inhibiting pro-inflammatory cytokines in neurons and what mechanism or transcription factors are involved during the process. Hence, to address these issues, we designed the experimental study with SH-SY5Y cells, as continuation to our previous study with pesticides aldicarb (ALD) and chlorpyrifos (CPF) to obtain some novel outcomes.

## MATERIALS AND METHODS

### Materials

Pesticides (ALD and CPF) and chemicals (Lithium lactate, NAD, 2, 4-dinitrophenyl-hydrazine reagent; DNPH, 1.0 N hydrochloric acid, Sodium Pyruvate, Ethylenediamine tetra-acetic acid; EDTA, 5-5’-dithiobis [2-nitrobenzoic acid]; DTNB, Trichloroacetic Acid; TCA, α-Ketoglutarate, Potassium ferrocyanide, Potassium cyanide, Succinate, Oxaloacetate, Cytochrome c, N-phenyl-p-phenylene diamine, agarose were commercially purchased from Sigma Aldrich, Takara Bio, Thermofisher Scientific etc.

### Cell culture reagents

SH-SY5Y cells were obtained from NCCS, Pune, India. Dulbecco’s modified eagle media (DMEM) and Ham’s F12 medium, fetal bovine serum (FBS), 1% penicillin-streptomycin were commercially purchased from Thermofisher scientific.

### Buffers

glycine, sodium phosphate, Tris-HCl (pH 7.2), Tris-HCl 0.1 M (pH 7.5), ptassium phosphate buffer (pH 7.4), 0.3 M Phosphate buffer (pH 7.6), dH_2_O were prepared in laboratory.

### Solutions

1 M NaOH 1 M, Guanidine hydrochloride 8 M, Ethanol/Ethyl Acetate (1:1 v/v), 0.1 M Trisodium isocitrate, 0.015 M Manganese chloride, 0.002 M Thiamine pyrophosphate, 0.25 M Potassium ferricyanide, 3 % BSA were purchased from Sigma Aldrich. All other reagents used were of analytical standard.

## Methods

### 1. Purification and characterization of human A2M

The A2M protein was isolated and purified from human blood plasma using ammonium sulfate precipitation followed by gel exclusion chromatography as per the method described in previous study (**Dixit et al., 2023**). A 5% (w/v) native PAGE was performed and the gel was stained with Coomassie brilliant blue R-250 (0.15% in 10% acetic acid). The gel was destained for 12 h in the destaining solution (10% acetic acid), and the purified A2M formed a single band on the gel.

### 2. Cell culture

The SH-SY5Y cell line was cultured in a medium containing 1:1 DMEM and Ham’s F12 medium, 10% FBS, and 1% penicillin-streptomycin (**Dixit et al., 2023**). The cells were treated with a standard solution of pesticides and α2M, accordingly, to perform the experiments. Cells were used at 3–7 passages. The cells were divided into five groups based on the treatment with pesticides and proteins to obtain results for various stress markers under standard conditions (37 C and 5% CO_2_). . Cells were marked under various groups:

***Group I:*** It was the control group comprising only SHSY5Y cells.

***Group II:*** It consisted of SH-SY5Y cells incubated with ALD and CPF (5 µM each) separately under standard conditions for 3 h.

***Group III:*** It comprised 2 sets of cells treated with pesticides ALD and CPF separately and each set subsequently treated with A2M (2 µM) under standard conditions for 3 h.

***Group IV:*** This group consisted of A2M alone treated SH-SY5Y cells for 3 h.

### 3. Analysis of pathophysiological marker of cell membrane damage; assay of lactate dehydrogenase (LDH)

LDH release is generally considered as marker of cellular damage and cell death upon acute or chronic injury in muscle and brain cells (**Kaja et al., 2015**). In the assay, the lactate is acted upon by lactate dehydrogenase to form pyruvate in the presence of NAD. The pyruvate forms pyruvate phenyl hydrazone with 2,4 dinitrophenyl hydrazine. The colour substrate developed is read at 420nm. The enzyme activity was expressed as IU/L (µmole pyruvate liberated/min/litre) and tissue enzyme activity was expressed as μmoles pyruvate liberated/min/mg protein (**Kumar et al., 2018**).

### 4. Analysis of neurotoxicty; acetylcholinesterase (AChE) assay

AChE activity is measured to assess exposure to neurotoxic compounds such as organophosphate and carbamate pesticides (**Lionetto et al., 2013**) in humans. It is a reliable biomarker of neurotoxicity attributed to its dose-dependent response to exposure to pollutants (**Bernal-Rey et al., 2020**). AChE enzymatic activity was assayed using standard method (**Sun et al., 2017**). AChE hydrolyses the acetylthiocholine to produce thiocholine and acetate. The liberated thiocholine reduces the dithiobis-nitrobenzoic acid (DTNB) liberating nitrobenzoate, which absorbs at 405 nm. The change in absorbance was recorded at 412 nm for 2 min at 30 sec interval using UV spectrophotometer and the activity was expressed as µmoles of acetylcholine iodide hydrolyzed /min/mg protein.

### 5. Analysis of protein carbonyls; major hallmark of oxidative damage

Oxidative stress is considered as sign of several neurodegenerative diseases, including Alzheimer’s, Parkinson’s, and Huntington’s disease (**Dixit et al., 2023**). Protein carbonyl levels are estimated to measure the extent of protein oxidation to predict the oxidative stress in early stages leading to neurodegeneration (**Dalle-Donne et al., 2006**). The measurement of protein carbonyls involve reaction of carbonyl group with 2,4-dinitrophenylhydrazine (DNPH), which leads to the formation of a stable 2,4-dinitrophenyl (DNP) hydrazone product (**Rogowska-Wrzesinska et al., 2014**). The optical density of each sample was read at 365 nm against the control. The level of protein carbonyls was expressed as nmoles /mg protein.

### 6. Analysis and estimation of mitochondrial enzymes

Mitochondrial TCA cycle is a primary source of cellular energy during which acetyl-CoA is oxidised to produce carbon dioxide, ATP and NADH. Various enzymes participate during this cycle to form intermediate substrates ultimately leading to the release of acetyl-CoA. Excessive free radicals generated by some toxicants damage the inner mitochondrial membrane, leading to compromised mitochondrial energy production and metabolism in the brain (**Vujic et al., 2021**). To assess the mitochondrial functioning, we performed the estimations of following TCA cycle enzymes-

#### a) Assay of Isocitrate dehydrogenase

The enzyme activity was assayed according to the standard method (**Kim et al., 2007**). Isocitrate dehydrogenase catalyzes the decarboxylative oxidation of threo-DS-isocitrate (2R-3S-isocitrate) by NADP+ or NAD+, yielding α-ketoglutarate, carbon dioxide, and NADPH or NADH. The reaction is magnesium/manganese dependent. Reduction of NADP+ /NAD+ to ADPH/NADH is read spectrophotometrically at 420 nm. The isocitrate dehydrogenase activity was expressed as nmoles of α-ketoglutarate liberated/hr/mg mitochondrial protein.

#### b) Assay of α-Ketoglutarate dehydrogenase

α-KGDH activity was determined from the rate of reduction of NAD+ in the presence of α -ketoglutarate (potassium salt). The colour substrate was measured at 460 nm. The activity of α-ketoglutarate dehydrogenase was expressed as nmoles of ferrocyanide liberated/hr/mg of mitochondrial protein (**Gibson et al., 2015**).

#### c) Assay of Succinate dehydrogenase

Succinate dehydrogenase catalyzes the oxidation of succinate to fumarate with the sequential reduction of enzyme-bound FAD and non-heme-iron. Reaction mixture consisted of 1 ml of phosphate buffer, 0.1 m1 of EDTA, 0.1 ml of BSA, 0.3 ml of sodium succinate, 40 mM sodium azide and 50 μM 2,6-dichloroindophenolate. This mixture was added to the cell lysate. The change in OD was recorded at 15 seconds interval for 5 min at 600 nm. The succinate dehydrogenase activity was expressed as nmoles of succinate oxidized/min/mg of mitochondrial protein (**Cimen et al., 2010**)

#### d) Assay of Malate dehydrogenase

Malate dehydrogenase is an oxidoreductase involving nicotinamide adenine dinucleotide the decrease in absorbance due to the oxidation of NADH is followed. Oxaloacetic acid reacts with NADH and H+ and releases Malic acid and NAD. Reaction was recorded at 340 nm. The enzyme activity was expressed as nmoles of NADH oxidized/min/mg of mitochondrial protein (**Mansouri et al., 2017**).

### 7. Analysis of mitochondrial dysfunction

Mitochondrial damage signifies the disruption of mitochondrial respiratory chain enzymes. Mitochondria are both producers as well as targets of ROS, which increases oxidative damage. Upon continuous damage, mitochondria lose their functional integrity and release more ROS molecules compromising neuronal functioning and accelerating neurodegenerative process (**Bishop et al., 2010**). To analyse mitochondrial dysfunction, biochemical assay of ATPase activity and NADH dehydrogenase was performed-

#### a) ATPase activity assay

Na+ / K+ ATPase activity is the process by which the sodium-potassium adenosine triphosphatase (Na/K-ATPase) enzyme transports ions across the cell membrane. NKA dysfunction has been linked to mitochondrial dysfunction in neurodegenerative diseases (**Zhang et al., 2022**). Na+ / K+ ATPase transports Na+ / K+ against concentration gradient at the cost of ATP molecule liberating an inorganic phosphate (Pi). Na+ / K+ ATPase activity was estimated from the amount of Pi liberated in the reaction. The blue colored substrate was read at 620 nm against blank in UV spectrophotometer. Enzyme activity was expressed as μmoles of phosphorus liberated/min/mg protein at 37°C (**Rule et al., 2016**).

#### b) Assay of NADH dehydrogenase

NADH dehydrogenase complex or NADH-ubiquinone oxireductase is a large, multisubunit protein complex that transfers electrons from NADH to ubiquinone (coenzyme Q1O) in the respiratory chain of mitochondrial complex I. To the reaction mixture containing phosphate buffer, potassium ferricyanide and NADH, cell mitochondrial lysate suspension was added. The change in OD was measured at 400 nm as function in time dependent manner (**Wang et al., 2016**). The activity of NADH dehydrogenase was expressed as nmoles of NADH oxidized/min/mg mitochondrial protein.

#### c) Assay of Cytochrome-c-oxidase

Cytochrome-c-oxidase is a terminal enzyme in the cellular respiration which reduces oxygen to water. This process creates an electrochemical gradient that powers the synthesis of adenosine triphosphate (ATP), which is used to fuel cellular processes. The activity of cytochrome-c-oxidase was assayed by the standard method in which the enzyme oxidizes the reagent (N-phenyl-p-phenylene diamine) to color end product (**Zhou et al., 2004**). The enzyme activity was expressed as nmoles of cytochrome oxidized /min/mg of mitochondrial protein.

### 8. Isolation and analysis of DNA fragmentation by agarose gel electrophoresis

DNA from SH-SY5Y cells after pesticides treatment and A2M treatment was isolated and extracted by using kit from Qiagen and following the manufacturer’s instruction. Sample DNA in microfuge tube was dissolved in TAE buffer (pH: 8.0) containing 40 % sucrose and a pinch of bromophenol blue marker dye. Agarose gel was prepared and the apparatus was set up equipped with the constant power supply of of 150 V. When the gel was run completely, the supply was removed and gel was then visualized by ethidium bromide (EtBr) in UV trans-illuminator (**Kašuba et al., 2017**).

### 9. Western blotting analysis

The nucleic components were isolated using a nucleic/cytosolic fractionation kit (Takara Bio) according to the manufacturer’s instructions. The protein content of each cellular extract was quantified by the Bradford assay. An equal amount of protein from each sample was separated by SDS-poly-acrylamide gel electrophoresis (10% gel) and transferred to the nitrocellulose membrane. The membranes were blocked with 5% skimmed milk and incubated with primary antibodies for TNF-α, NF-ĸβ, p53, iNOS, IL-1β, Caspase-3, Caspase 9, anti Bax and anti-Bcl-2 antibody (Santa-Cruz Scientific, USA) and HRP-conjugated secondary antibody (Thermo Fisher, USA) followed by enhanced chemiluminescence detection (Thermo scientific, USA) (**Kim et al., 2008**). The nuclear fraction was incubated with anti-Nrf2 and anti-β actin antibody (Santa-Cruz Scientific, USA) for normalization of positive loading control. Relative band intensities were quantified using Image analysis software.

### 10. Statistical analysis

All the grouped data was evaluated using SPSS software. Hypothesis testing method included one-way analysis of variance (ANOVA), followed by least significant difference (LSD) test P<0.05 indicates statistical significance. All the results were expressed as mean ± S.D for triplicates in each group.

## RESULTS

### 1) Purification of A2M

**Fig. 2** shows the results from the purification of human A2M from human plasma. Native gel electrophoresis of human A2M consisted of several fractions to rule out the purest fraction containing A2M. In this figure, lane 1 consist of human blood plasma, lane 2 consisting of 20-40% ammonium sulfate fraction and lane 3 has purified A2M (180 KDa), obtained after gel filtration chromatography.

**Figure 2:**
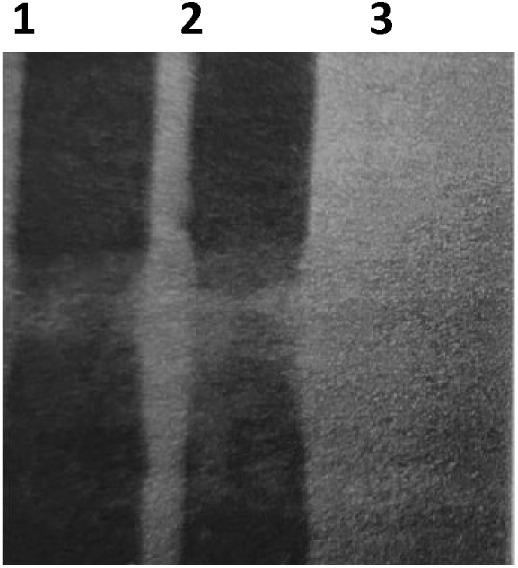
Native gel electrophoresis of human A2M consisting of the purified A2M (lane 3).

### 2) Effect of A2M and ALD/CPF on pathophysiological marker enzyme LDH in control and experimental group of SH-SY5Y cells

Table 1 & **fig. 3** show the effect of ALD, CPF and A2M on Lactate dehydrogensae (LDH); a pathophysiological marker enzymes in control and experimental group of SH-SY5Y cells. The results indicated that the levels of LDH were significantly (P<0.05) increased in group II when compared with control cells (group I). However, supplementation of A2M to pesticides treated cells significantly lowered the LDH levels (group III) as compared to group II. A2M alone treated cells did not show any significant change in the enzyme’s activity (group IV).

**Figure 3:**
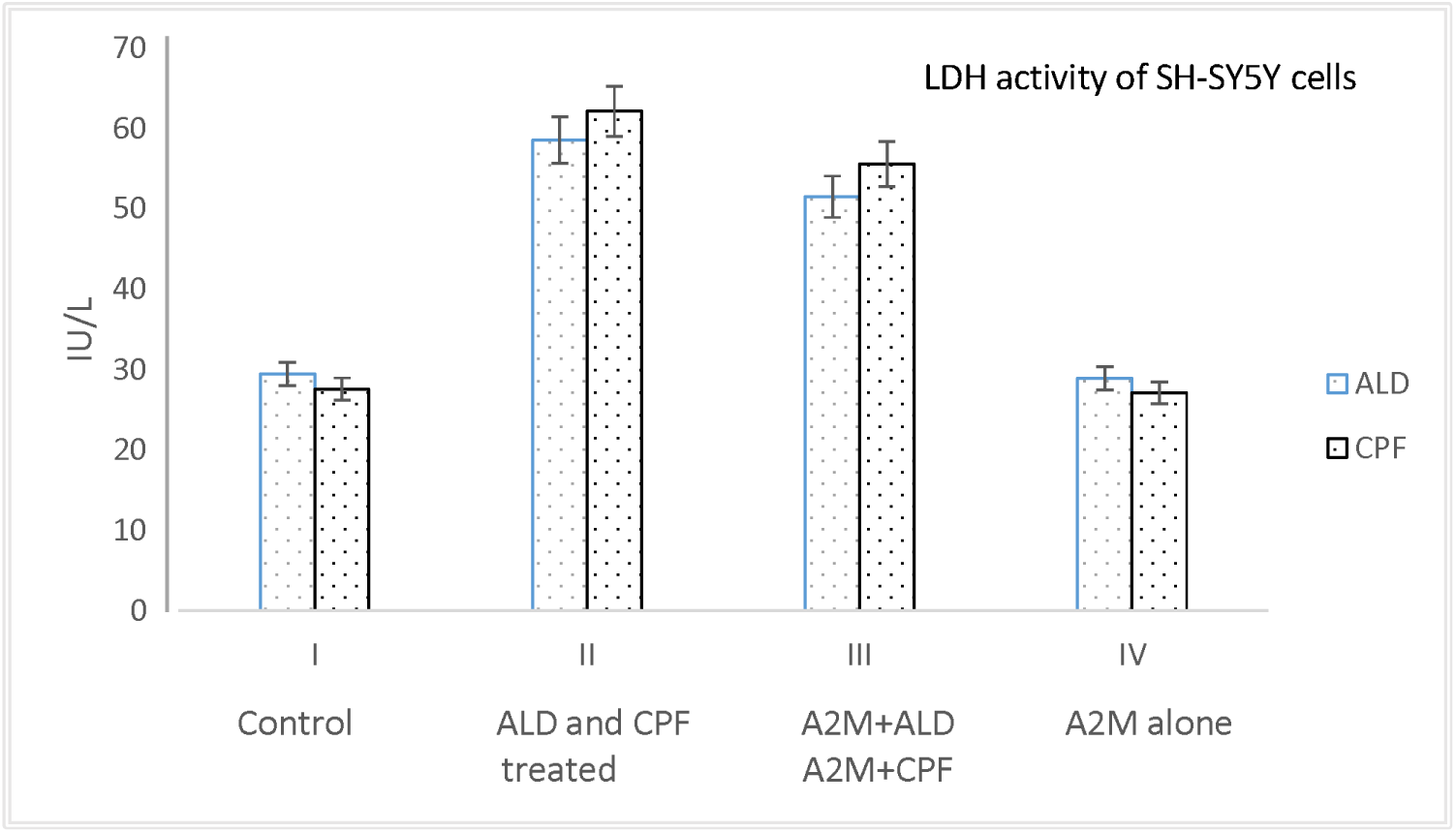
LDH: Lactate dehydrogenase. Activity is expressed as IU/L. Results are expressed as mean + SD for different sets of experiments (3 set/enzyme/group for ALD treated and A2M induced cells, 3 set/enzyme/group for CPF treated and A2M induced cells)

**Table 1:** Effect of pesticides ALD and CPF and treatment with A2M on pathophysiological marker LDH on control and experimental group of SH-SY5Y cells. Group I represents control, Group II represents ALD treated cells and CPF treated cells, Group III represents treatment of pesticides induced cells with A2M and group IV represents A2M treated cells alone.

| Groups | ALD | CPF |
| --- | --- | --- |
| I | 29.44±3.51 | 27.57±2.22 |
| II | 58.53 ± 4.21 | 62.13±5.73 |
| III | 51.49±2.33 | 55.55±3.88 |
| IV | 27.13±3.61 | 23.62± 2.14 |

### 3) Effect of A2M and ALD/CPF on the activities of AChE in SH-SY5Y cell homogenate of control and experimental groups

The effects of ALD, CPF and A2M on the activity of acetyl cholinesterase (AChE) in control and experimental group of SH-SY5Y are shown in Table 2 and **fig. 4**. The obtained results showed that a significant (p<0.05) inhibition in the activity of AChE in ALD and CPF treated cells (group II) compared with control (group I). However, upon simultaneous treatment of A2M with ALD and CPF was able to preserve the AChE activity levels (group III). The present results suggest the healthy role of A2M in significantly countering the harmful effects of pesticides ALD and CPF. A2M alone treated cells did not cause any significant alteration on the AChE activity (group IV) compared to control.

**Figure 4:**
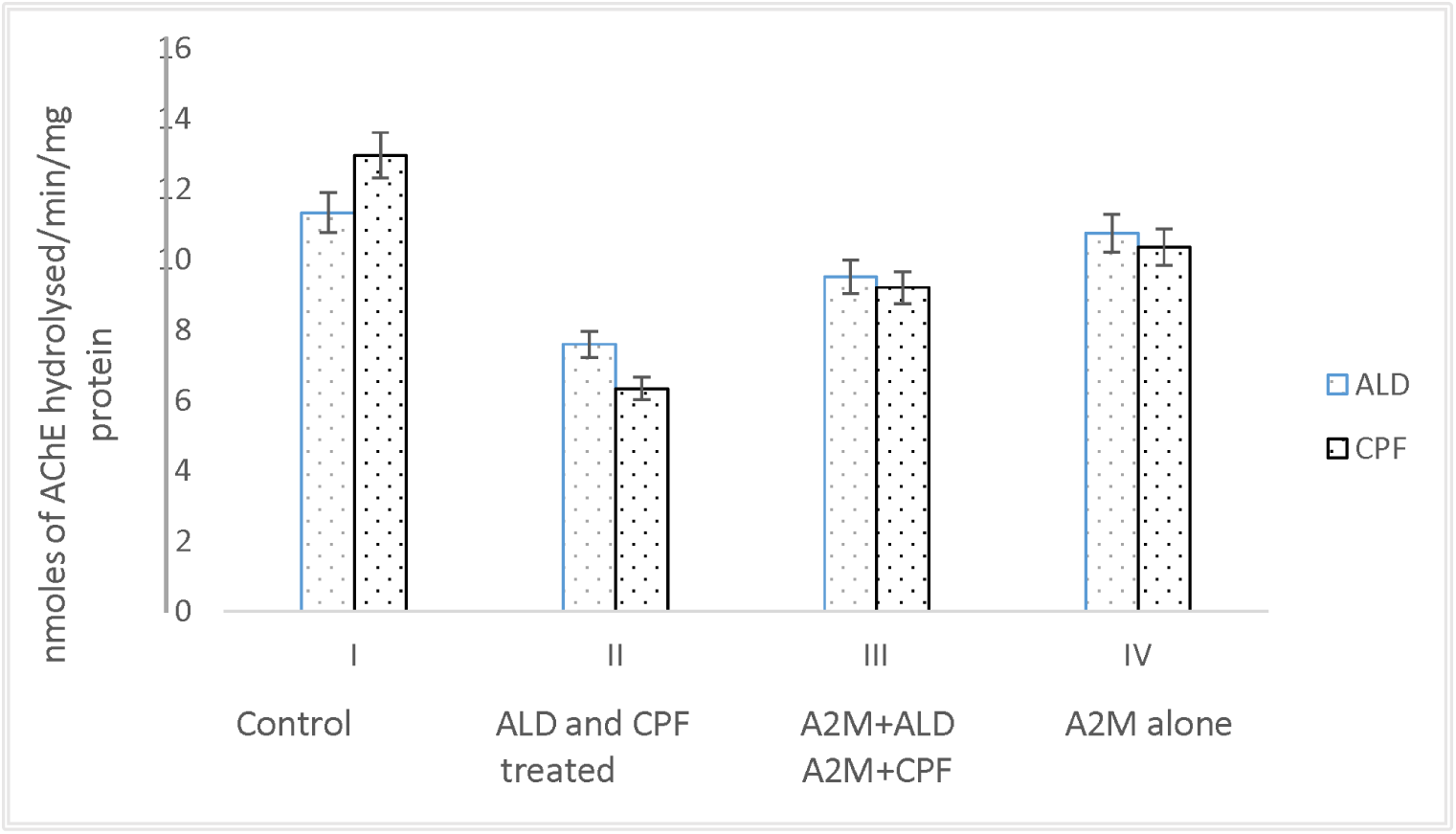
Results are expressed as mean + SD for different sets of experiments (3 set/ enzyme/group for ALD treated and A2M induced cells, 3 set/enzyme/group for CPF treated and A2M induced cells)

**Table 2:** Effect of pesticides ALD and CPF and treatment with A2M on AChE levels in control and experimental group of SH-SY5Y cells. Group I represents control, Group II represents ALD treated cells and CPF treated cells, Group III represents treatment of pesticides induced cells with A2M and group IV represents A2M treated cells alone.

| Groups | ALD | CPF |
| --- | --- | --- |
| I | 11.33±0.94 | 12.96±0.55 |
| II | 7.59 ± 0.81 | 6.34±0.46 |
| III | 9.51±0.35 | 9.20±0.68 |
| IV | 10.75±0.72 | 10.36± 0.59 |

### 4) Effect of A2M and ALD/CPF on the levels of protein carbonyl in mitochondrial lysate of control and experimental group of SH-SY5Y cells

Pesticides act as pro-oxidants and triggers oxidative stress in multiple organs. Table 3 & **fig. 5** illustrates the effects of ALD, CPF and A2M on the protein oxidation in control and experimental group of SH-SY5Y cells. Protein carbonyl (PC) is the marker for protein oxidation. There was a significantly increased PC (p<0.05) contents was shown in ALD and CPF treated SH-SY5Y cells (group II) as compared with control group (group I), confirming the susceptibility of neuronal cells to oxidative stress. However, administration of A2M to ALD and CPF treated cells, A2M significantly decreased (p<0.05) the PC level (group III) compared with ALD and CPF treated cells. No significant difference was observed in cells treated with A2M alone (group IV) when compared to control cells (group I).

**Figure 5:**
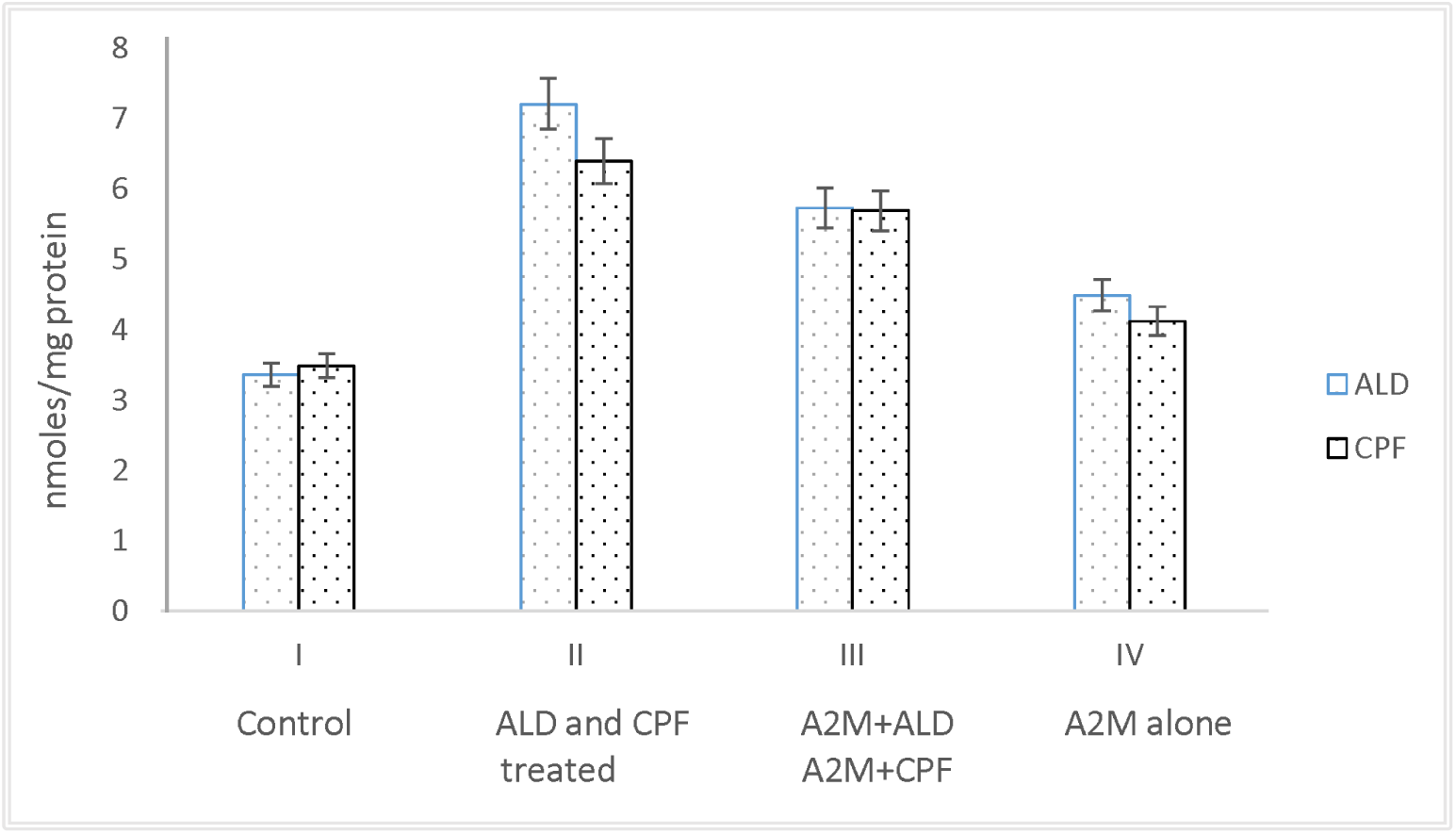
Results are expressed as mean + SD for different sets of experiments (3 set/ enzyme/group for ALD treated and A2M induced cells, 3 set/enzyme/group for CPF treated and A2M induced cells)

**Table 3:** Effect of pesticides ALD and CPF and treatment with A2M on protein carbonyls in control and experimental group of SH-SY5Y cells. Group I represents control, Group II represents ALD treated cells and CPF treated cells, Group III represents treatment of pesticides induced cells with A2M and group IV represents A2M treated cells alone.

| Groups | ALD | CPF |
| --- | --- | --- |
| I | 3.36±0.15 | 3.49±0.16 |
| II | 7.21 ± 0.55 | 6.40±0.66 |
| III | 5.73±0.27 | 5.69±0.35 |
| IV | 4.49±0.22 | 4.12± 0.08 |

### 5) Analysis and estimation of ALD, CPF and A2M on the activities of mitochondrial TCA cycle enzymes in control and experimental group of SH-SY5Y cells

The effects of pesticides ALD and CPF treated control and experimental groups of SH-SY5Y cells and their further treatment with A2M on mitochondrial TCA cycle enzymes and complex I and III are shown in Table 4. The results indicated that the activities of mitochondrial enzymes were significantly (P<0.05) decreased in pesticides induced cells (group II), compared with the control cells (group I). However, treatment with A2M in combination with the pesticides alleviated their negative effects and significantly increased (P<0.05) the activities of the mitochondrial enzymes (group III). A2M alone treated cells did not show any significant changes in the mitochondrial enzyme activities (group IV).

**Table 4:**
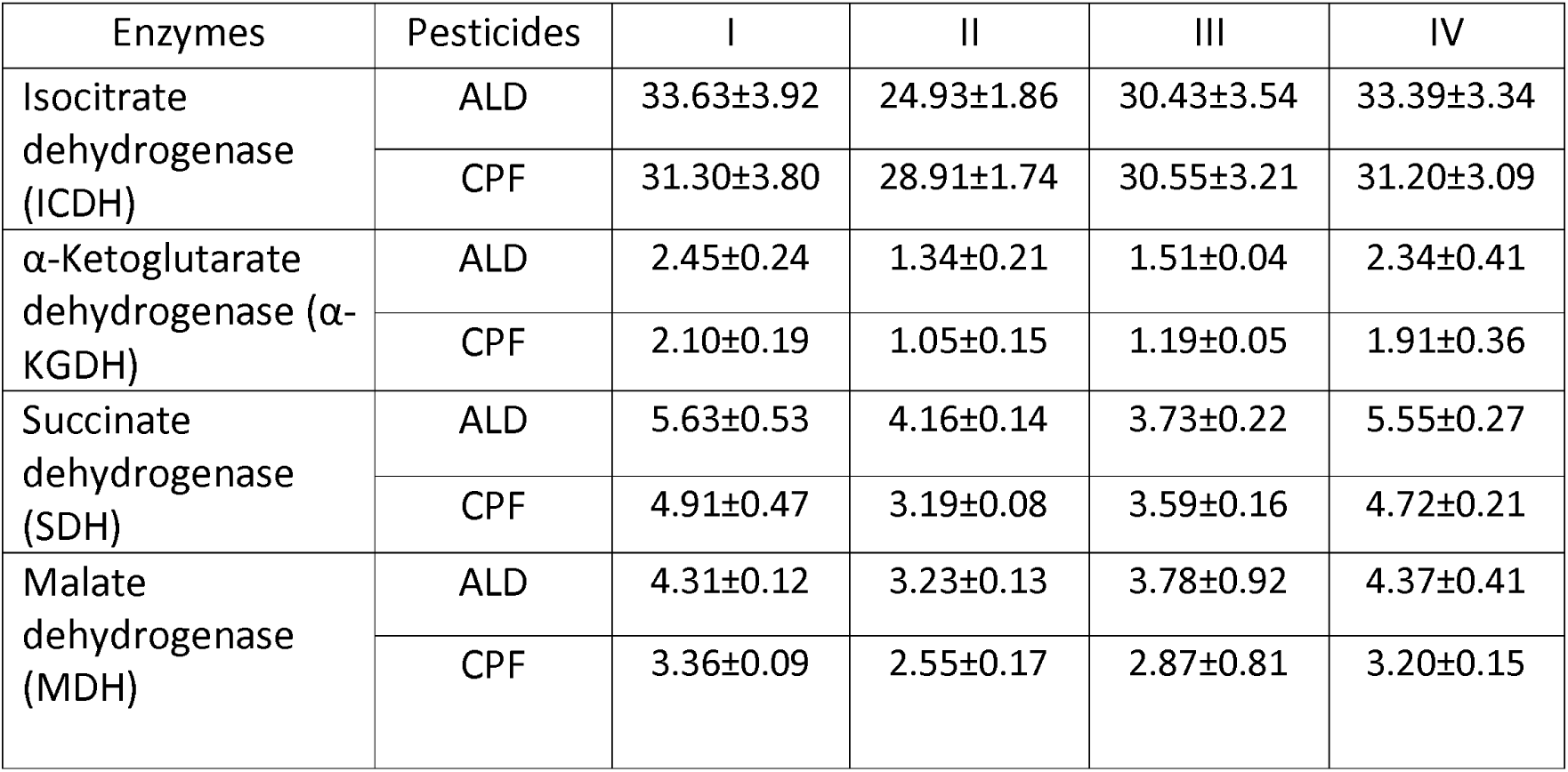
Effect of pesticides ALD and CPF and treatment with A2M on mitochondrial TCA cycle enzymes in control and experimental group of SH-SY5Y cells. Group I represents control, Group II represents ALD treated cells and CPF treated cells, Group III represents treatment of pesticides induced cells with A2M and group IV represents A2M treated cells alone.

Units: ICDH-nmoles of ketoglutarate formed/hr/mg protein; α-KGDH - nmoles of ferrocyanide formed/hr/mg protein; SDH - nmoles of succinate oxidized/min/mg protein; MDH -nmoles of NADH oxidized/min/mg protein.

Values are expressed as mean + SD for different sets of experiments (3 set/ enzyme/group for ALD treated and A2M induced cells, 3 set/enzyme/group for CPF treated and A2M induced cells) with (P<0.05) as statistical significance.

### 6) Analysis of mitochondrial dysfunction in control and experimental group of SH-SY5Y cells

**Fig. 6** represent the activity of Na+ / K+ ATPase pump in response to the ALD, CPF and A2M treated group in control and experimental group of SH-SY5Y cells. Na+ /K+ ATPase activity was significantly (P<0.05) decreased in ALD and CPF induced rats (group II) suggesting the impairment of ATPase activity could be one of the underlying biochemical mechanism that leading to early cognitive CNS dysfunctions. However, administration of A2M with ALD and CPF treated cells, was able to prevent the inhibition of this enzyme pump (group III), thereby indicating that A2M protect ATPase from pesticides induced depletion of its activity. A2M alone treated cells did not cause any significant alteration on the ATPase activity (group IV) compared to control group of SH-SY5Y cells. The effect of pesticides ALD and CPF treated SH-SY cells for complex I (NADH dehydrogenase) and complex III (Cytochrome-c-oxidase) and the effect of A2M, are given in table 5.

**Figure 6:**
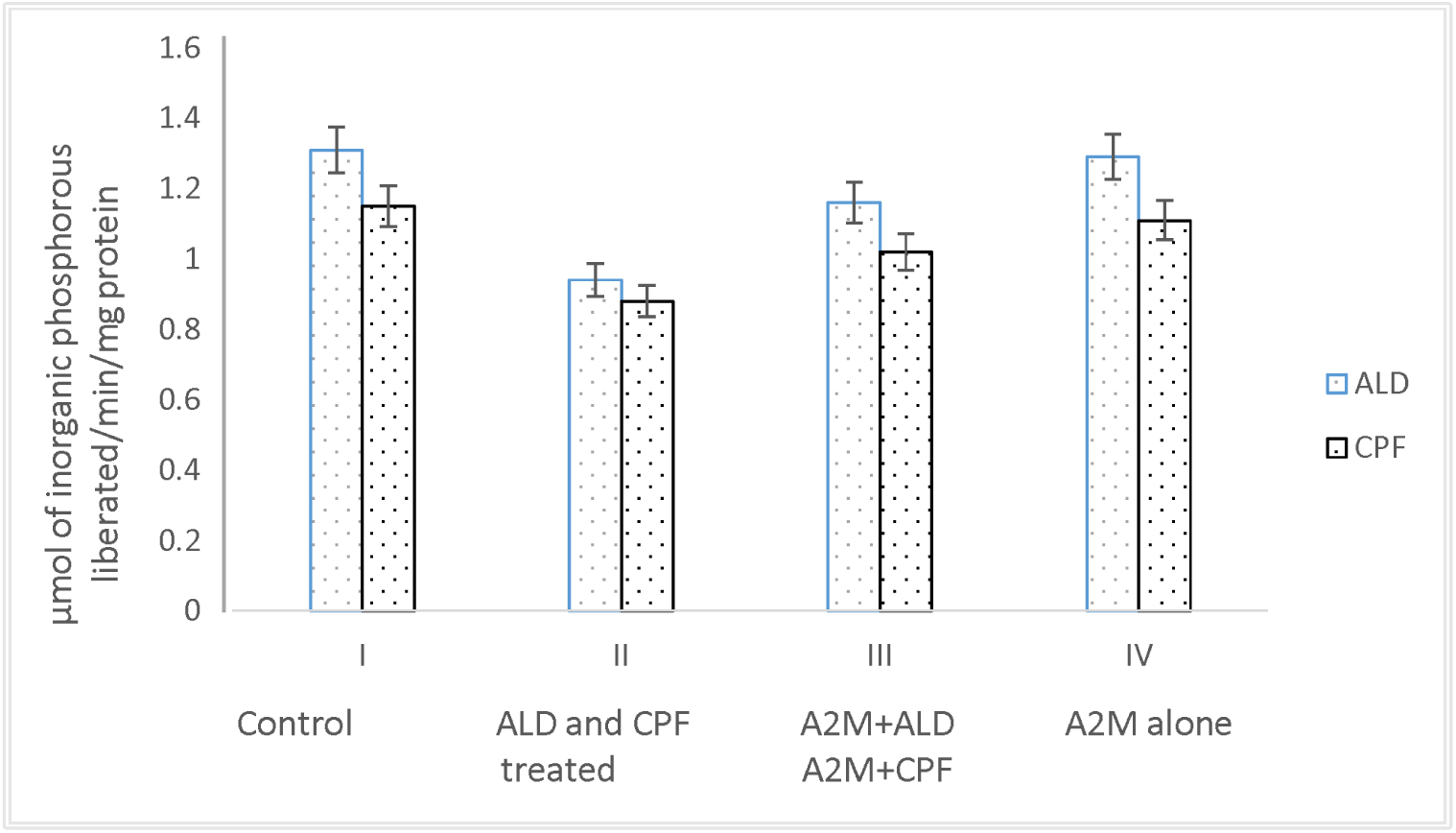
Results are expressed as mean + SD for different sets of experiments (3 set/ enzyme/group for ALD treated and A2M induced cells, 3 set/enzyme/group for CPF treated and A2M induced cells)

**Table 5:**
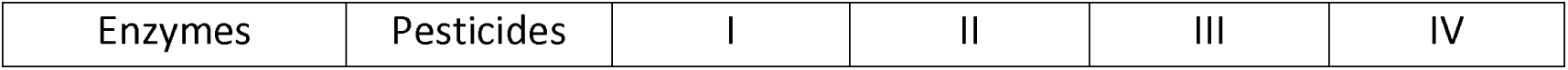
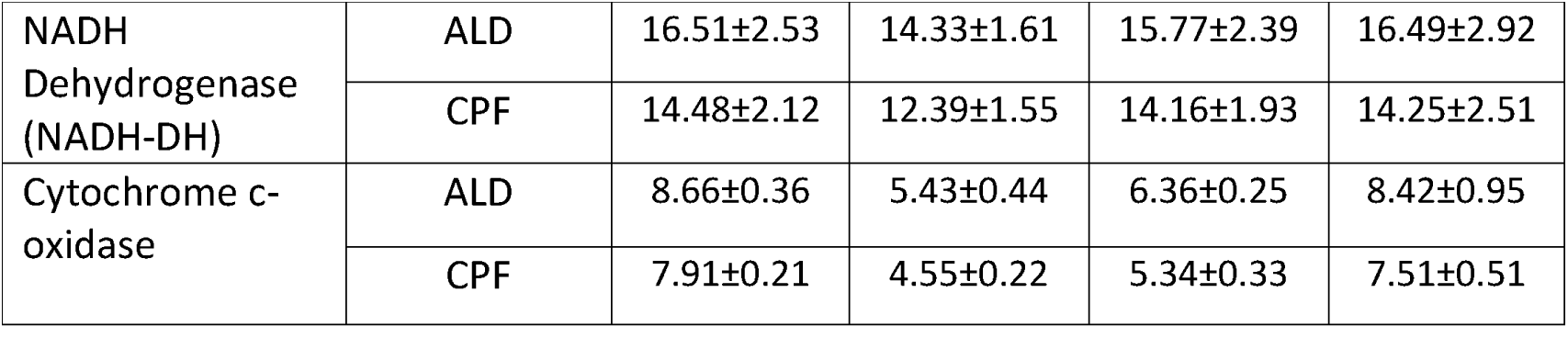
Effect of pesticides ALD and CPF and treatment with A2M on mitochondrial dysfunction of complex I NADH Dehydrogenase and complex III cytochrome-c-oxidase in control and experimental group of SH-SY5Y cells. Group I represents control, Group II represents ALD treated cells and CPF treated cells, Group III represents treatment of pesticides induced cells with A2M and group IV represents A2M treated cells alone.

| Enzymes | Pesticides | I | II | III | IV |
| --- | --- | --- | --- | --- | --- |
| NADH Dehydrogenase (NADH-DH) | ALD | 16.51±2.53 | 14.33±1.61 | 15.77±2.39 | 16.49±2.92 |
|  | CPF | 14.48±2.12 | 12.39±1.55 | 14.16±1.93 | 14.25±2.51 |
| Cytochrome c-oxidase | ALD | 8.66±0.36 | 5.43±0.44 | 6.36±0.25 | 8.42±0.95 |
|  | CPF | 7.91±0.21 | 4.55±0.22 | 5.34±0.33 | 7.51±0.51 |

Units in which the values are expressed are as follows:

NADH-DH - nmoles of NADH oxidized/min/mg protein Cytochrome-c-oxidase - nmoles of cytochrome/min/mg protein.

Values are expressed as mean + SD for different sets of experiments (3 set/ enzyme/group for ALD treated and A2M induced cells, 3 set/enzyme/group for CPF treated and A2M induced cells) with (P<0.05) as statistical significance.

### 7) Effect of A2M and ALD/CPF on DNA fragmentation of SH-SY5Y cells

DNA pattern analysis of SH-SY5Y cells treated with pesticides ALD and CPF (lane 2) and exposed to A2M (lane 3) later is shown in **fig. 7**, along with the control group of cells (group I, lane 1) and A2M induced cells alone (lane 4). This agarose gel electrophoresis was performed to assess the DNA damage in neuronal cells after being introduced by the pesticides and to investigate the role of A2M. Pesticides induced cells (group II, lane 2) gave smear like pattern suggesting DNA contamination or DNA degradation (highlighted in lane 2). However, when the group II was treated with A2M, the band in lane 3 (group III) showed prevented DNA smear and fragmentation as compared to lane 2. Lane 1 and Lane 4 (group IV) showed no fragmentation.

**Figure 7:**
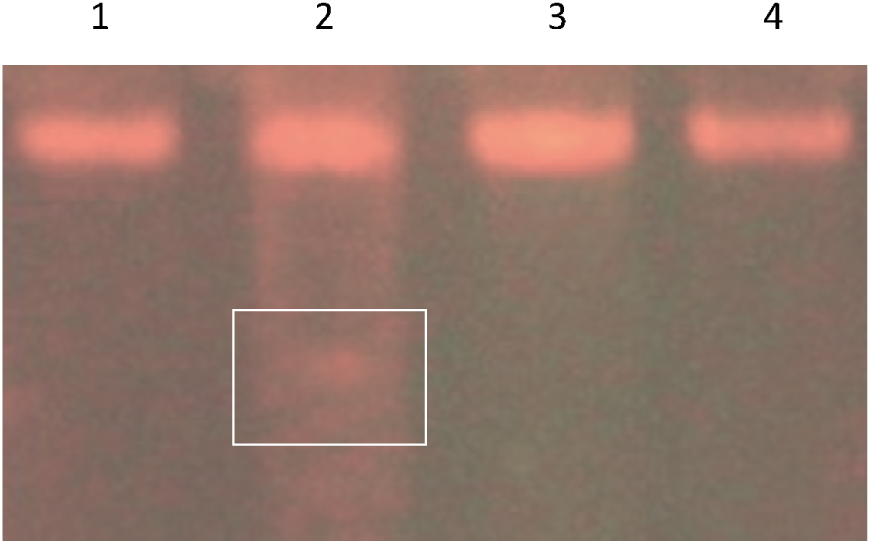
DNA agarose gel electrophoresis results of control and experimental groups of SH-SY5Y cells. Lane 1-SH-SY5Y cells, Lane 2-SH-SY5Y cells + ALD/CPF, Lane 3-SH-SY5Y cells+ ALD/CPF + A2M, Lane 4-SH-SY5Y cells+ A2M

### 8) Effect of pesticides and A2M on expression of inflammatory marker proteins in control and experimental group of SH-SY5Y cells

A2M binds with cytokines such as TNF-α, NF-ĸβ and IL-1β (**Dixit et al., 2023**). These cytokines promote neurotoxicity and neuroinflammation by silencing cell survival mechanisms, promoting Fas signals, enhancing glutamate levels from microglia, increasing Ca+2 permeability in NMDA receptors etc. (**Sun et al., 2023**). A2M traps and neutralize these cytokines and decrease the inflammation in various cells (**Dixit et al., 2022b**). In **fig. 8a**, western blot expressions of these inflammatory markers are shown in control and experimental groups of SH-SY5Y cells.

**Figure 8:**
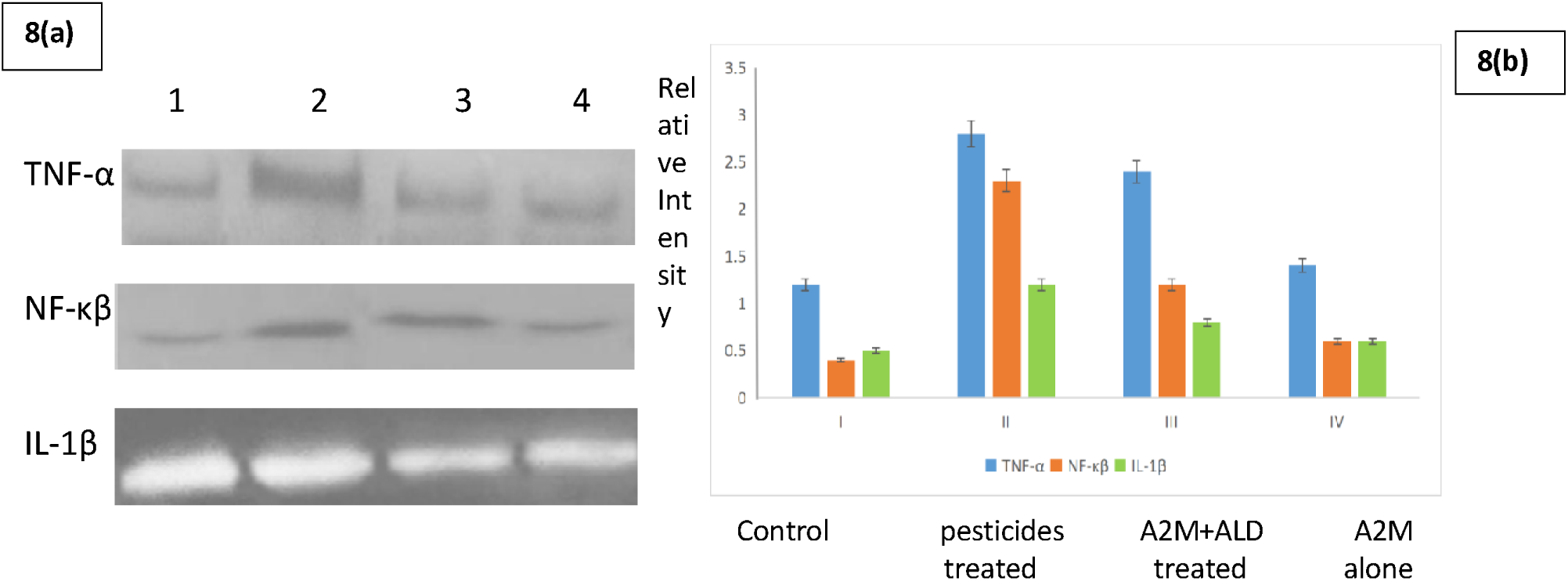
8(a): Western blot expression of inflammatory marker proteins in control and experimental group of SH-SY5Y cells. Lane 1-SH-SY5Y cells, Lane 2-SH-SY5Y cells + ALD/CPF, Lane 3-SH-SY5Y cells+ ALD/CPF + A2M, Lane 4-SH-SY5Y cells+ A2M. **8(b)**: Relative intensity of the proteins expressed in control and experimental groups of SH-SY5Y cells.

Pesticides induced neuronal SH-SY5Y cells (group I) showed increased expression of these markers (lane 2) as compared to control cells (group II) (lane 1). Lane 3 shows SH-SY5Y cells induced with pesticides and treated with A2M (group III) showed decreased expression of these inflammatory markers as compared their Lane 2 expressions. A2M treated cells (lane 4) didn’t show any significant expression of these inflammatory markers *(positive loading control* β*-actin was used but not shown)*. **Fig. 8b** represents the data expressing the respective protein levels which were quantified using the image analysis software and results are expressed in relative intensity. The values represent relative levels of protein expression.

### 9) Effect of pesticides and A2M on Bax, Bcl-2, Caspase-3, Caspase-9 ; apoptotic protein expressions in control and experimental group of SH-SY5Y cells

The mitochondria-dependent pathway is one of the most important pathways of apoptosis, which is regulated by the B cell lymphoma-2 (e.g.,Bcl-2) family proteins, including pro-apoptotic proteins (Bax) and anti-apoptotic proteins (e.g., Bcl-2). Caspase-3 is a proteolytic enzyme that regulate other caspases (e.g., Caspase-3), cause DNA damage and alter cell’s morphology to induce apoptosis during cancer progression and neuronal degeneration (**Chota et al., 2021**).

There is a small amount of knowledge regarding the expression patterns and role of A2M in apoptosis. So, in our study, results in **fig.9** suggests the positive role of A2M in regulating the apoptotic marker proteins. In **fig. 9a**, western blot expressions of these apoptotic markers proteins (Bax, Bcl-2, Caspase-3, Caspase-9) are shown in control and experimental groups of SH-SY5Y cells. Pesticides induced neuronal SH-SY5Y cells (group II) showed increased expression of these markers (lane 2) as compared to control cells (lane 1), but in Bcl-2 expression, pesticides were shown to reduce its expression *(positive loading control* β*-actin was used but not shown)*.

**Figure 9:**
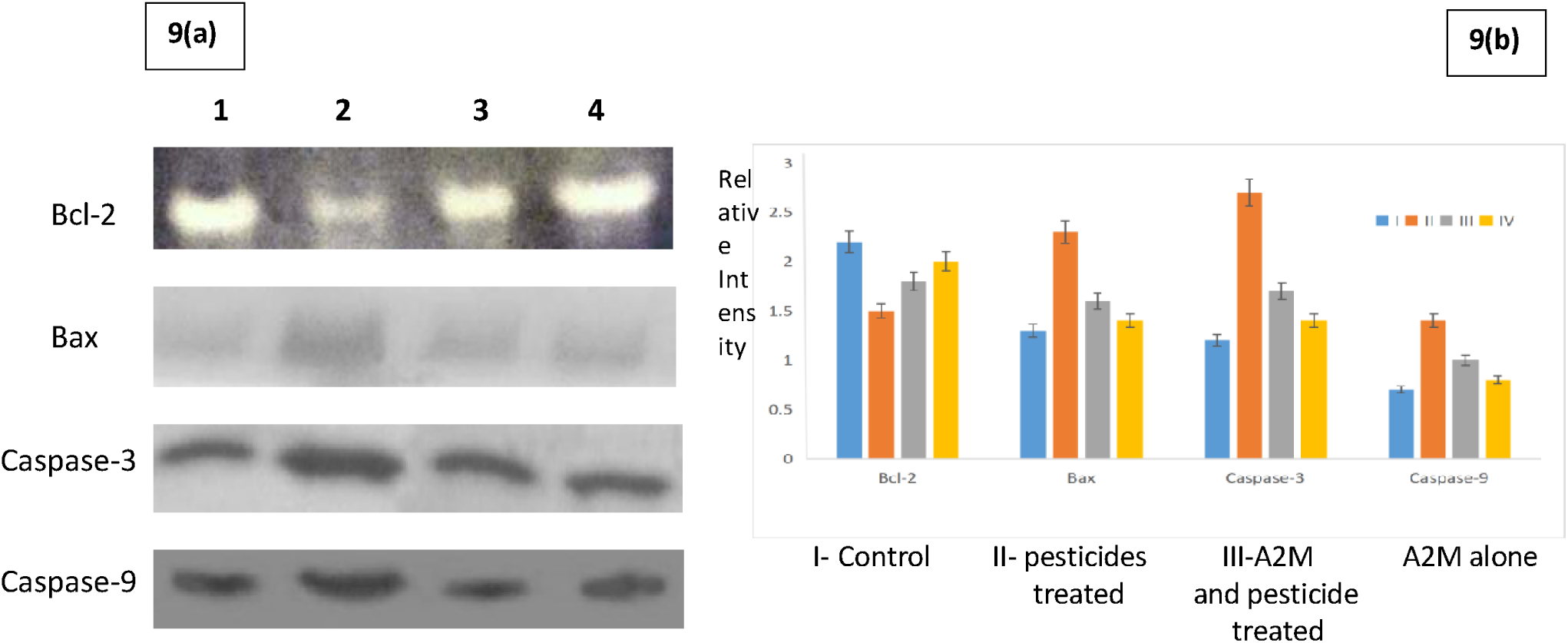
**9(a)**: Western blot expression of apoptotic marker proteins in control and experimental group of SH-SY5Y cells. Lane 1-SH-SY5Y cells, Lane 2-SH-SY5Y cells + ALD/CPF, Lane 3-SH-SY5Y cells+ ALD/CPF + A2M, Lane 4-SH-SY5Y cells+ A2M. **9(b)**: Relative intensity of the proteins expressed in control and experimental groups of SH-SY5Y cells.

Lane 3 shows SH-SY5Y cells induced with pesticides and treated with A2M (group III) showed decreased expression of these apoptotic markers as compared their Lane 2 expressions, while in Bcl-2 expression profile, A2M was found to restore Bcl-2 expression. A2M treated cells (lane 4) didn’t show any significant expression of these inflammatory markers. **Fig. 9b** represents the data expressing the respective protein levels which were quantified using the image analysis software and results are expressed in relative intensity. The values represent relative levels of protein expression.

### 10) Effect of pesticides and A2M on Nrf2 expression in control and experimental group of SH-SY5Y cells

Nrf2 is a transcription factor that regulates cellular redox balance and inflammation. It coordinates cellular defense processes against the pathological hallmarks of neurodegeneration, such as oxidative stress, neuroinflammation, mitochondrial dysfunction, and protein aggregation. Oxidative stress is associated with neuronal cell death during the pathogenesis of neurodegenerative diseases. There is a limited yet ongoing research shows that A2M helps in regulating Nrf2/KEAP1 pathway in necrosis (**Fang et al., 2024**). So we decided to investigate A2M’s function in Nrf2 pathway during pesticides induced neurotoxicity in neuronal cells.

Western blot analyses of Nrf2 in cytosolic and nucleus fraction of the control and experimental group of SH-SY5Y cells is shown in **fig. 10**. Pesticides induced group II showed slight reduction in expression of nuclear and cytosolic Nrf2 (lane 2) when compared to the control (lane 1). However, when group II was treated with A2M (group III), it resulted in significant increased expression of Nrf2 in its cytosolic and nuclear fractions as compared to the observations in group II, suggesting its role in balancing and regulating Nrf2 activity during pesticides induced neurotoxicity in neuronal SH-SY5Y cells. In group IV, A2M treatment alone didn’t have any noticeable observation in Nrf2 expression pattern(lane 4). β-actin was used as positive loading control.

**Figure 10:**
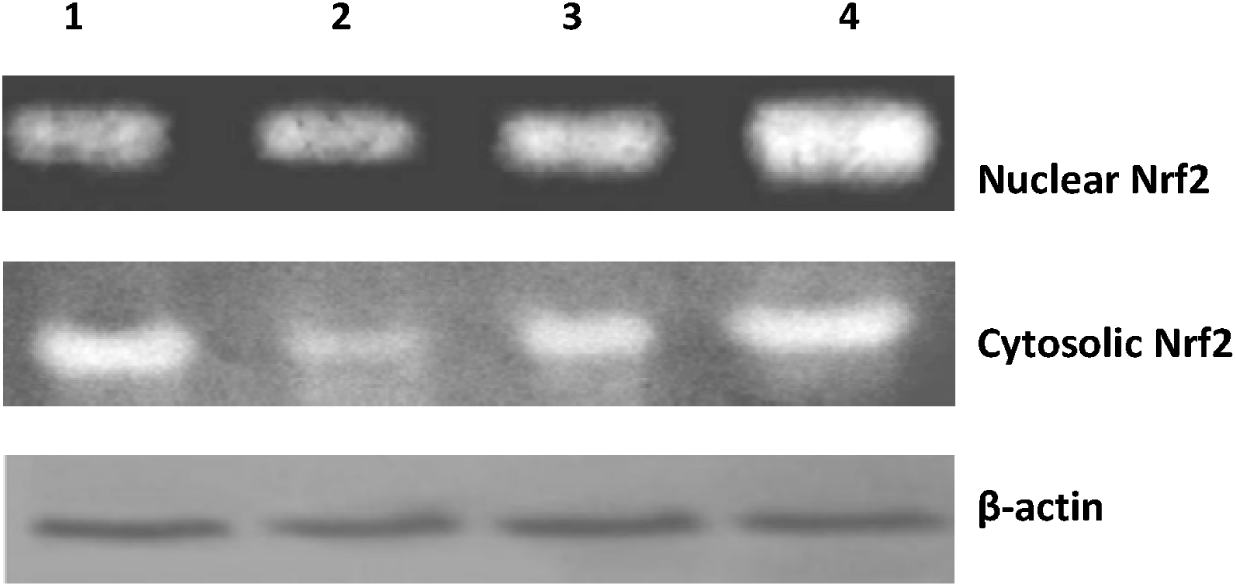
Western blot expression of Nrf2 nuclear and cytosolic fraction in control and experimental group of SH-SY5Y cells.

## DISCUSSION

Pesticides such as organophosphates and carbamates have found to provoke multiple deleterious and harmful effects in human body but most of them operate via the production of oxidative stress damage (**Dixit et al., 2023**). They accumulate in cells and over time, they start producing ROS and damage biomolecules such as lipids, proteins and DNA, thereby interfering with the biochemical processes ultimately leading to inflammation induced apoptotic cell death in living beings (**Dixit et al., 2022b**). Long term exposure to pesticides can cause inflammation in neuronal cells and the brain, which can lead to neurodegeneration. They activate glial cells, astrocytes and microglia, which produce pro-inflammatory factors such as the tumor necrosis factor-α (TNF-α), interleukin-1β (IL-1β), interleukin-1β (IL-10), nitric oxide (NO), and cyclooxygenase-2 (COX-2) that further damage neurons. This leads to neuroinflammation which leads to development of neurodegenerative diseases like Alzheimer’s and Parkinson’s (**Hossain et al., 2017**).

Various studies have shown the role of neuroinflammatory markers in the occurrence, diagnosis, and treatment of neurodegenerative diseases (**Rauf et al., 2022**). Alpha-2 macroglobulin (A2M) is a serum protein in the human circulatory system that act as guardian during inflammatory cellular injury by trapping and functionally inhibiting proteases (**Dixit et al., 2022a**). It also binds several cytokines, including interleukin (IL)-6, platelet-derived growth factor (PDGF), nerve-growth factor, tumor-necrosis factor (TNF)-α, and IL-1β, making it a crucial component during the interactions between several cytokines and the process of inflammation, by decreasing these inflammatory mediators (**Dixit et al., 2023**).

Lactate dehydrogenase (LDH) is a cytosolic enzyme that is released into the medium when cells are damaged through apoptosis, necrosis, or other events that cause membrane integrity decline. Its proliferation in the serum is an indicator of cellular damage (**Kaja et al., 2015**). In our findings, we demonstrated that ALD and CPF administrated SH-SY5Y cells provoked a marked elevation in the enzyme LDH activity LDH activities indicating neuronal cellular damage. Administration of A2M attenuated ALD and CPF induced neuronal toxicity as revealed by the decreased levels LDH activity suggests A2M imparting cellular protection against pesticides induced toxicity in SH-SY5Y cells by stabilizing the cellular membrane against the damage induced by these toxicants.

Acetylcholinesterase or AChE is an enzyme which hydrolyses the acetylcholine (ACh), a neurotransmitter. It is widely distributed throughout the body mainly in cholinergic nerves and erythrocytes (**Kalafatakis et al., 2015**). AChE activity is a standard biomarker of pesticide poisoning. Organophosphates and carbamates have been demonstrated to cause a decrease in activities of AChE in erythrocytes and CNS of living organisms (**Lionetto et al., 2013**).

In this present study, ALD and CPF exposure to SH-SY5Y cells resulted in a significant decrease in AChE levels. Probing the mechanism, which is likely to be inhibition of AChE’s active sites by these pesticides, and thereby causing a significant decline in its activity leading to progression of neurodegenerative disorders. A2M treatment significantly suppressing the pesticides toxicity in AChE activity suggests its neuroprotective potential, which could hint at its interaction with AChE or other molecular factors which regulate AChE activity.

Tissue proteins are vulnerable to ROS levels. ROS modifies the amino acids such as proline, arginine, lysine, and histidine which produce carbonyl moieties as a result of, protein oxidation and can be used as early parameter of protein damage (**Luceri et al., 2017**). Alteration of different intracellular proteins including major enzymes and structural proteins in the neuron of brain leads to the neurofibrillary degeneration (**Iqbal and Grundke-Iqbal, 2008**). PCs are suggested among major contributor for the loss of cellular function under oxidative stress in brain and age-related neurodegenerative disorders (**Bueno et al., 2010**).

Results demonstrate the increase in protein carbonyls (PCs) contents after administration of ALD and CPF in SH-SY5Y cells indicate an increased oxidative modification protein content in brain, increasing the chances of neurotoxicity. On A2M treatment, the PC content was significantly declined. Our data suggest that attenuating action of A2M is due to the inhibition of free radical induced protein oxidation and that this effect is mediated by endogenous activation of certain antioxidants.

Mitochondrial damage defines the prolonged mitochondrial exposure to ROS species and resulting in their loss of functional integrity combined with the production of more ROS molecules compromising neuronal functioning and accelerating neurodegenerative process (**Bishop et al., 2010**). Experimental findings shows the administration of pesticides ALD and CPF caused impairment in mitochondrial enzyme complex activity as indicated by decrease in the activity levels of (complex-I) NADH dehydrogenase (NADH-DH), (complex-II) Succinate dehydrogenase (SDH), (complex-III) Cytochrome-C-oxidase, Isocitrate dehydrogenase (ICDH), α-Ketoglutarate dehydrogenase (KGDH) and Malate dehydrogenase (MDH). These enzyme complexes are involved in ATP production by the process of oxidative phosphorylation in mitochondria. In pesticides induced mitochondrial injury, free radicals and oxidative stress are major implication, consequently inducing ROS toxicity interferring with oxidative phosphorylation and the TCA cycle in mitochondria, preventing mitochondrial ATP production (**Panov et al., 2007**).

Decrease in the concentration of complex III cytochrome-c-oxidase leads to a decrease in the uptake of oxygen, resulting in low respiratory rate. Thus, reduction in mitochondrial cytochrome content, results in a loss of activities of oxidative phosphorylation capacity. ALD and CPF induced decrease in the activities of complex I NADH dehydrogenase depletes the reducing equivalents like NADH and NADPH, which are normally utilized during the production of reduced glutathione to counter oxidative damage of mitochondrial components (**Karthikeyan et al., 2007**). Results confers that A2M supplementation significantly attenuated the pesticides induced mitochondrial TCA cycle enzyme damage and mitochondrial dysfunction exerting its antioxidant and neuroprotective effect at mitochondrial level.

Na+ /K+ ATPase is a membrane bound lipid dependent enzyme which catalyzes the active transport of Na+ and K+ in CNS in order to regulate cellular homeostasis and the ionic concentration gradient for neuronal excitability (**Hussien et al., 2013**). When neurons become toxic, intracellular ATP levels are rapidly declined and Na+ -K+ -ATPase activity decreases which could signal for neural apoptosis (**Ijima et al., 2006**). Pesticides have shown to have strong affinity of interaction with cellular membrane lipids by altering structural and functional integrity of cell membrane and also effects membrane bound enzymes, such as Na+ /K+ ATPase (**Zbarsky et al., 2005**).

In this study, ALD and CPF exposure prevent the normal functioning of Na+ /K+ ATPase activity. The decrease in cellular ATP levels due to mitochondrial dysfunction leads to inhibition of the Na+ /K+ across the neuronal synaptic membrane whose cascade leads to the degeneration of axons in numerous neurodegenerative disorders (**Millecamps and Julien, 2013**). While supplementation of A2M with pesticides ALD and CPF, A2M was found to restore the activities of Na+ -K+ -ATPase strengthening the integrity of cell membrane thereby ensuring the normal physiological functions of brain cells.

Free radicals mediated oxidative damage to DNA results in the functional and structural alterations in the DNA leading to proliferative cell death. DNA fragmentation is considered as a marker and usual characteristic feature of apoptosis cell death (**Rong et al., 2021**). In this study, treatment of pesticides significantly increased DNA damage. This could be attributed to the hydrophobic and lipophilic nature of pesticides which is stored in various cells and tissues and triggers ROS production and DNA damage (**Abdel-Daim et al., 2013**). Our findings suggests the pesticides induce mitochondrial dysfunction activating some proteases which led to DNA fragmentation and apoptosis. A2M has pivotal role as protease inhibitor, hence treatment with A2M reduced the DNA fragmentation, by grappling the proteases releases during mitochondrial depletion.

Inflammatory cytokines such as TNF-α, NF-ĸβ, IL-1β, IL-6 etc. play an important role in the pathogenesis of oxidative stress induced neurodegenerative disorders such as Parkinson’s disease and Alzheimer’s disease. A2M levels increase during inflammatory diseases and neutralize pro-inflammatory cytokines by its bait and trap mechanism.

In the present study, we observed that carbamate pesticide (ALD) and organophosphate (CPF) increased the expression of cytokines (TNF-α, NF-ĸβ, IL-1β) in SH-SY5Y cells as shown in their western blot profile. Pesticides causes cellular damage, oxidative stress and free radicals that activates the transcription of multiple inflammatory genes. Our study shows A2M restoring the neuroinflammation by reducing the levels of inflammatory cytokines, possibly indicating its therapeutic role in early prognosis in inflammation induced injury in neurons (**Lagrange et al., 2022**).

B-cell lymphoma 2 (Bcl-2) is a family of proteins which govern the mitochondria-dependent pathway for apoptosis. Among them, Bcl-2-associated X protein (Bax) and Bcl-2 have been well studied and examined in apoptotic cascades (**Anilkumar and Prehn, 2014**) . Caspases are the cysteine proteases that execute programmed cell death. Caspases-2, -3, and -9 are some of the caspases that trigger apoptosis (**Papaliagkas et al., 2007**). Pesticides have now been understood to stimulate the transcription and translation of apoptotic proteins such as Bax and p53 but downregulation of Bcl-2 protein, which prevents apoptosis and promote cell survival (**Nur et al., 2022**).

Findings suggest that pesticides triggered increased expression of apoptotic (Caspase-3, Caspase-9) and pro apoptotic (Bax) proteins, and were found to reduce the expression of anti apoptotic Bcl-2 protein. A2M treatment to the pesticides treated apoptotic proteins (Bax, Caspase-3, Caspase-9) was found to lower their expression, owing to its inhibition tendency for proteolytic nature of these proteins. However, in case of Bcl-2 expression, where pesticides reduced its expression level, A2M restored its activity, suggesting its bait and trap mechanism regulated inhibition of pesticides ALD and CPF or hinting at its involvement in ER stress pathway in neurons, which is also a mechanism of pesticides induced neurotoxicity (**Chen et al., 2023**).

Nuclear factor erythroid 2 related factor -2 (Nrf2) is a transcription factor which regulates cell survival against neurodegenerative diseases, aging, diabetes,inflammation and cancer. NRF2 balances the expression of antioxidant proteins, detoxification enzymes, and other cytoprotective genes which protects cells from oxidative stress (**Ma, 2013**). Nrf2 activation modulates neuronal inflammatory responses and its mechanism is widely studied to develop therapeutic and preventive strategies for the management of inflammation mediated neurodegenerative disorders. A2M regulates oxidative stress response by activating the Keap1/Nrf2 signaling pathway, which is a key defense mechanism in necrosis (**Fang et al., 2024**).

In the present study, we have investigated the potential molecular mechanisms underlying the protective effect of A2M in pesticides induced neurodegeneration via regulation/activation of Nrf2-mediated gene expression pathways. Normally, Nrf2 is located in the cytoplasm in its inactive form. When oxidative response triggers, it migrates to the nucleus where it activates and stimulate the gene expression of Nrf2-regulated antioxidant response elements (ARE) which imparts protection to neuron against oxidative stress (**Bergström et al., 2011**). As pesticides impart toxicity by triggering the levels of reactive oxygen species, they subsequently hinders the functioning of Nrf2, due to imbalance in the antioxidant system of cell, as observed from the experimental findings in **fig. 10**. A2M treatment to the pesticides treated cells restored the functioning of Nrf2 both in cytosolic and nucleus fractions, corroborating with its functional relationship with Nrf2 protein. Mechanism behind this suggests the involvement of the unique bait sequence of A2M which is believed to interact with upregulation of other transcription factors to further enhance the activation of Nrf2 in nucleus during oxidative response.

## CONCLUSION

This study elucidates the mechanisms underlying pesticide (ALD and CPF)-induced neuronal toxicity, emphasizing their role in triggering mitochondrial-mediated apoptosis and subsequent inflammatory apoptotic cell death in SH-SY5Y cells. Based on our previous study, we continued the investigation of A2M in mitochondrial enzymes and its relationship with inflammatory and apoptotic marker proteins, in response to pesticide mediated neuronal injury.The investigation highlights the pivotal role of Alpha-2-Macroglobulin (A2M) in counteracting these adverse effects. A2M effectively mitigates oxidative stress in mitochondria, enhances cellular antioxidant defense systems, facilitates DNA repair, and suppresses neuronal inflammation and apoptosis. A schematic conclusion of the study is given below in the **fig. 11**.

**Figure 11:**
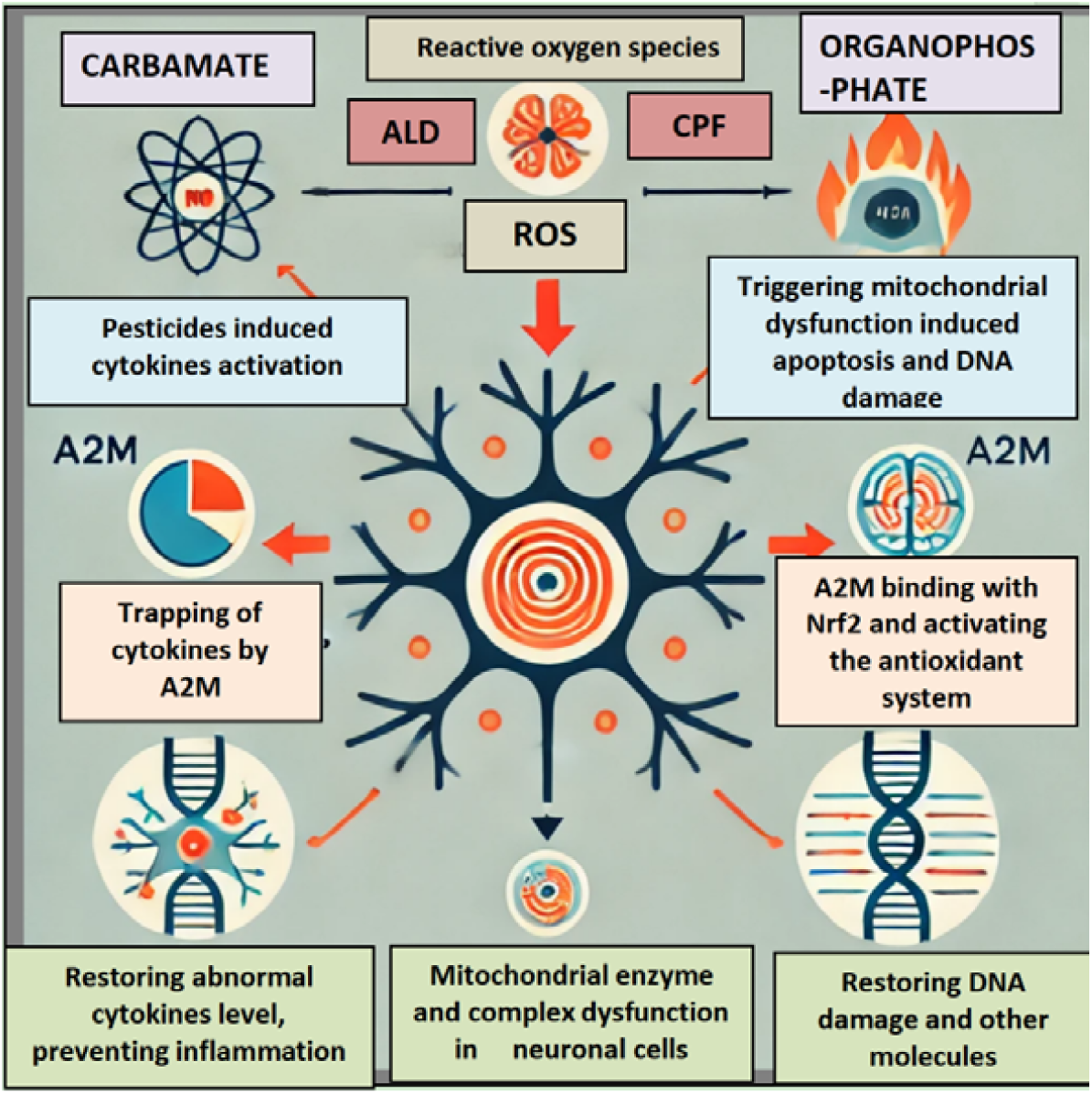
Graphical representation of highlighting A2M’s role in mitigating oxidative stress in mitochondria by enhances cellular antioxidant defense systems, facilitating DNA repair and suppressing of neuronal inflammation and apoptosis.

This protective action preserves neuronal integrity and functionality, reducing the likelihood of pesticide-induced neurodegeneration. Importantly, this research provides the first evidence of A2M’s interaction with apoptotic markers and mitochondrial enzyme complexes, offering novel insights into its therapeutic potential for combating pesticide-mediated neuronal injury during preclinical stage. However, further studies are needed to get more insights into nuclear morphological assessment.

## ACKNOWLEDGEMENTS

Author is thankful to Indian Council of Medical Research (ICMR), New Delhi, India for the Senior Research Fellowship grant and instrumentation centre facility of National Institute of Pathology, ICMR, New Delhi, India.

## DATA AVAILABILITY STATEMENT

The data supporting the findings of this study are available within the article and its supporting materials.

